# Trade-Offs Between Hepatic Host Defense and Metabolic Programs Underlie Sex-Biased Diseases

**DOI:** 10.1101/2022.01.07.475423

**Authors:** Joni Nikkanen, Yew Ann Leong, William C. Krause, Denis Dermadi, J. Alan Maschek, Tyler Van Ry, James E. Cox, Ethan J. Weiss, Omer Gokcumen, Ajay Chawla, Holly A. Ingraham

## Abstract

Current concepts in evolutionary medicine propose that trade-offs and mismatches with a shifting environment increase disease risk. While biological sex also impacts disease prevalence, contributions of environmental pressures to sex-biased diseases remain unexplored. Here, we show that sex-dependent hepatic programs confer a robust (~300%) survival advantage for male mice during lethal bacterial infection. The transcription factor BCL6, which masculinizes hepatic gene expression at puberty, is essential for this advantage. However, protection by BCL6 comes at a cost following dietary excess, resulting in overt fatty liver and glucose intolerance in males. Deleting hepatic BCL6 reverses these phenotypes but markedly lowers male fitness during infection, thus establishing a sex-dependent tradeoff between host defense and metabolic systems. We suggest that these tradeoffs, coupled with current environmental pressures, drive metabolic disease in males.

---

Infections are one of the strongest evolutionary pressures shaping human physiology and disease. As such, the immune system and host defense responses are often prioritized at the expense of other physiological systems (1, 2). As a result, disease-causing genetic variants may be maintained in the population if they simultaneously improve survival during infection. For example, variants in human *G6PD* and *APOL1* increase the risk of sickle cell anemia and chronic kidney disease (3) but exert a strong protective effect against malaria and trypanosome infections, respectively. These studies highlight the notion of an ‘evolutionary trade-off’ whereby natural selection fails to optimize two traits simultaneously, causing increased fitness for one trait at the expense of another, ultimately elevating disease risk.

Shifting environments also magnify disease risk associated with trade-offs, resulting in so-called mismatch diseases (4). Thus, the initial benefit of a trait becomes detrimental in a new environment. For example, a mismatch between the modern diet and our genetic legacy is proposed to account for the high prevalence of chronic metabolic diseases, such as type 2 diabetes (T2D), heart disease, and fatty liver (5). However, although the evolutionary mismatch theory can explain chronic diseases affecting immunity and metabolism, their marked sex bias in the human population is poorly understood. Notably, men carry a much higher disease burden for common metabolic disorders compared to premenopausal women (6–8). Similarly, survival outcomes for some infectious diseases exhibit a strong sex bias (9). Together, inherent sex differences in physiological systems dictate disease progression in males and females. Here, we examined the relationship between biological sex during a dietary challenge and infection.

Prior studies found that mice housed at thermoneutral temperature (30°C) are susceptible to the metabolic consequences of chronic dietary excess (10, 11) and infection (12). We, therefore, used thermoneutral conditions to examine the relationship between metabolism and host defense mechanisms in C57BL/6J males and females (Fig. 1A). Despite an equivalent increase in body weight and fat mass when fed a high-fat diet (HFD, Fig. 1B, C), only male mice develop severe fatty liver and overt macrosteatosis (Fig. 1D-F and fig. S1A). Using these same housing conditions, we then asked how sex affects host fitness during infection with a sublethal dose of *E. coli* (strain O111:B4). Males are far less susceptible to infection, showing a strikingly higher survival rate and greater body mass preservation than females (Fig. 1G and fig. S1B). Spleen bacterial counts were equivalent in both sexes, indicating no differences in pathogen clearance (Fig. 1H). Greater fitness in males was also observed following activation of host immunity by the endotoxin lipopolysaccharide (LPS) (Fig. 1I). Collectively, our results expose a stark relationship, specifically in males, between hepatic fat accumulation following dietary excess and host fitness following bacterial infection.

**Figure 1.**
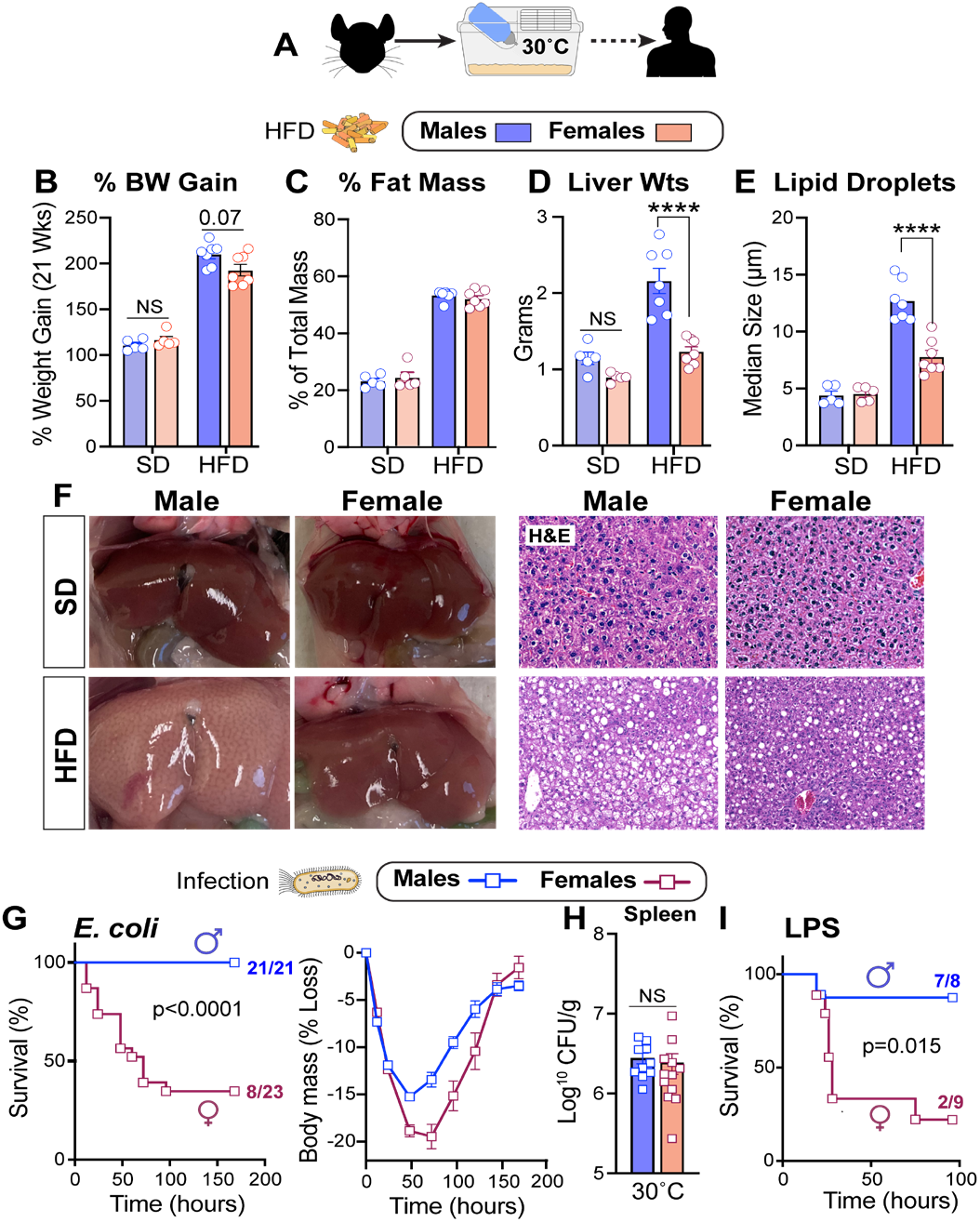
HFD and Infection Elicit Strong Sex-Dependent Phenotypes in Mice. **(A)** Schematic of housing conditions. **(B)** Body weight gain, **(C)** fat percentage, **(D)** liver weights, **(E)** hepatic lipid droplet size, and **(F)** whole livers with corresponding H&E staining after 21 weeks of SD or HFD. **(G)** Survival curves and body weights (n=9-10 per sex) of C57BL/6J mice infected with E. coli (1×10^8^ CFU). Data pooled from two experiments and analyzed by two-way ANOVA. **(H)** Bacterial CFUs of mice infected with *E. coli.* (I) Survival curves of mice treated with LPS (2 mg/kg). All mice were housed at 30°C. Data are presented as mean ± SEM, NS: not significant, ****p < 0.0001.

In searching for a sex-dependent hepatic factor that might mediate these divergent outcomes, the transcriptional repressor B-cell lymphoma six protein (BCL6) emerged as a top candidate given its role in hepatic lipid handling (13, 14) and its enrichment in the male liver (15). Indeed, *Bcl6* transcripts and protein are highly expressed in male hepatocytes and livers (Fig. 2A, 2B). Conditional deletion of *Bcl6* in the liver (*Bcl6^AlbCre^*) (Fig. 2B) feminizes the adult male liver and eliminates their male-biased gene signature (Fig. 2C, D, Data S1, and fig. S2A). Profiling active enhancers and promoters for acetylated histone three lysine 27 (H3K27ac) by ChIP-Seq also revealed an essential role of BCL6 in maintaining sex-dependent hepatic chromatin acetylation and male-biased H3K27ac peaks (Fig. 2C, D, and fig. S2B).

**Figure 2.**
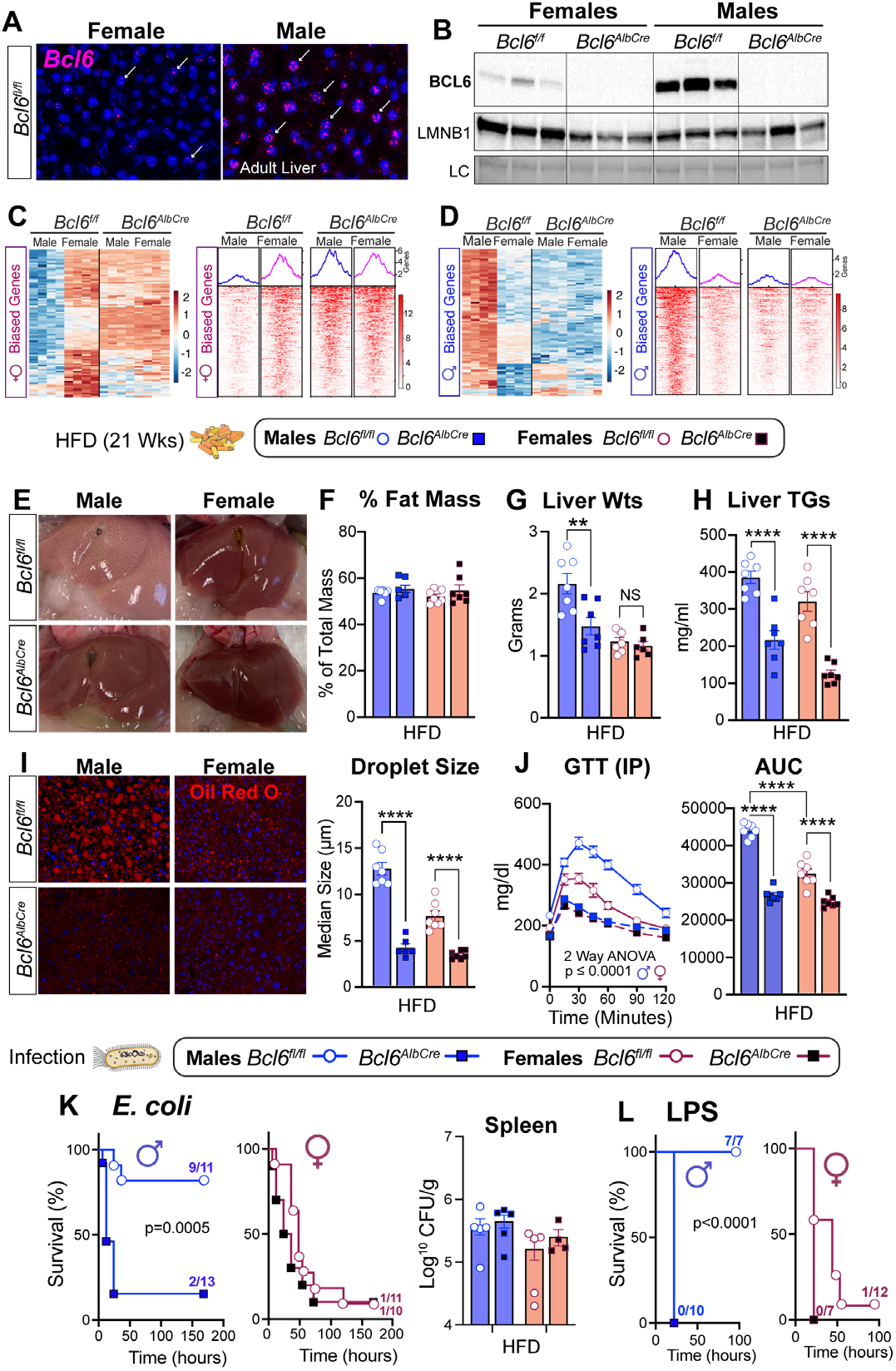
BCL6 Is Essential for Maintaining Hepatic Maleness and Survival to Infection But Impairs Metabolism. **(A)** In situ hybridization for *Bcl6* (magenta, white arrows) in livers of mice housed at 22°C. **(B)** Immunoblotting for BCL6 and LMNB1 in liver nuclear extracts of 8-week-old *Bcl6^f/f^* and *Bcl6^AlbCre^* mice housed at 22°C. **(C)-(D)** Heatmaps for top 100 female/male-bi-ased genes (filtered by fold change) with all female/male-biased H3K27ac peaks (q-value for both <0.05). **(E)** Livers, **(F)** fat per-centage, **(G)** liver weights, **(H)** liver triglycerides of mice fed HFD at 30°C. **(I)** Hepatic fluorescent labeling (ORO) and quantification of lipid droplet size (red) from mice fed HFD. Nuclei stained with DAPI (blue). **(J)** Glucose levels and AUC following GTT in mice fed HFD for 8 weeks. **(K)** Survival curves and bacterial counts of mice infected with E. coli (1×10^8^ CFU). **(L)** Survival curves of mice treated with LPS (1.75 mg/kg). Data are presented as mean ± SEM, NS: not significant. **p<0.01; ****p < 0.0001.

Having established the masculinizing role of BCL6 in hepatic gene signatures, we asked if BCL6 is essential for maintaining the distinct sex-specific outcomes to HFD and infection. Indeed, deleting hepatic *Bcl6* abolished all morphological hallmarks of fatty liver in males without changing their total fat mass or percent body weight gain (Fig. 2E, F and fig. S2C, D). Liver weights, hepatic triglycerides (TAGs), lipid accumulation, and droplet size were all significantly reduced in *Bcl6^AlbCre^* male mice (Fig. 2G-I), consistent with a prior study that found BCL6 blocks the breakdown of fat by lowering fatty acid oxidation (14). Hepatic TAGs also fell in *Bcl6^AlbCre^* mice fed a standard diet (SD, fig. S2E). Although more subtle, eliminating the low amounts of BCL6 present in female livers attenuated hepatic TAGs (Fig. 2G-I). Loss of hepatic BCL6 markedly improved glucose homeostasis in mutant male cohorts fed either HFD or SD and abolished any notable sex differences in this metabolic parameter (Fig. 2J and fig. S2F). In stark contrast to the improved metabolic state in *Bcl6^AlbCre^* males, their survival dropped precipitously after E. coli infection or LPS-treatment, plummeting to levels exhibited by control females (Fig. 2K, L). Pathogen clearance was unaffected in *Bcl6A^lbCre^* mice (Fig. 2K). Thus, high hepatic BCL6 in males is essential for optimizing host fitness during infection but drives fatty liver and glucose intolerance during dietary excess, reinforcing the strong inverse association between lipid handling and host defense responses.

Low survival in females is closely correlated with extremely high plasma TAGs leading to hyperlipidemia (Fig. 3A) – a known marker of poor survival during human sepsis (16). Likewise, compromised survival in infected *Bcl6^AlbCre^* males was linked with a substantial rise in levels of circulating TAGs species, similar to those of infected females (Fig. 3B, C, and fig. S3A). We then asked if high plasma TAGs contribute directly to poor survival in females by leveraging ANGPTL4 KO mice (17) that clear out TAGs due to increased lipoprotein lipase (LPL) activity (Fig. 3D). Normalizing triglycerides in infected *Angptl4*^-/-^ females restored both survival and body weights (Fig. 3E and fig. S3B). *Angptl4*^-/-^ males also showed a comparable drop in TAGs and remained resistant to infection (fig. S3C, D). Conversely, increasing plasma TAGs using poloxamer 407 (P407), a synthetic inhibitor of LPL, worsened survival of males following infection (Fig. 3F). Our results establish that the marked sex differences in infection outcomes are tightly linked with levels of circulating lipids.

**Figure 3.**
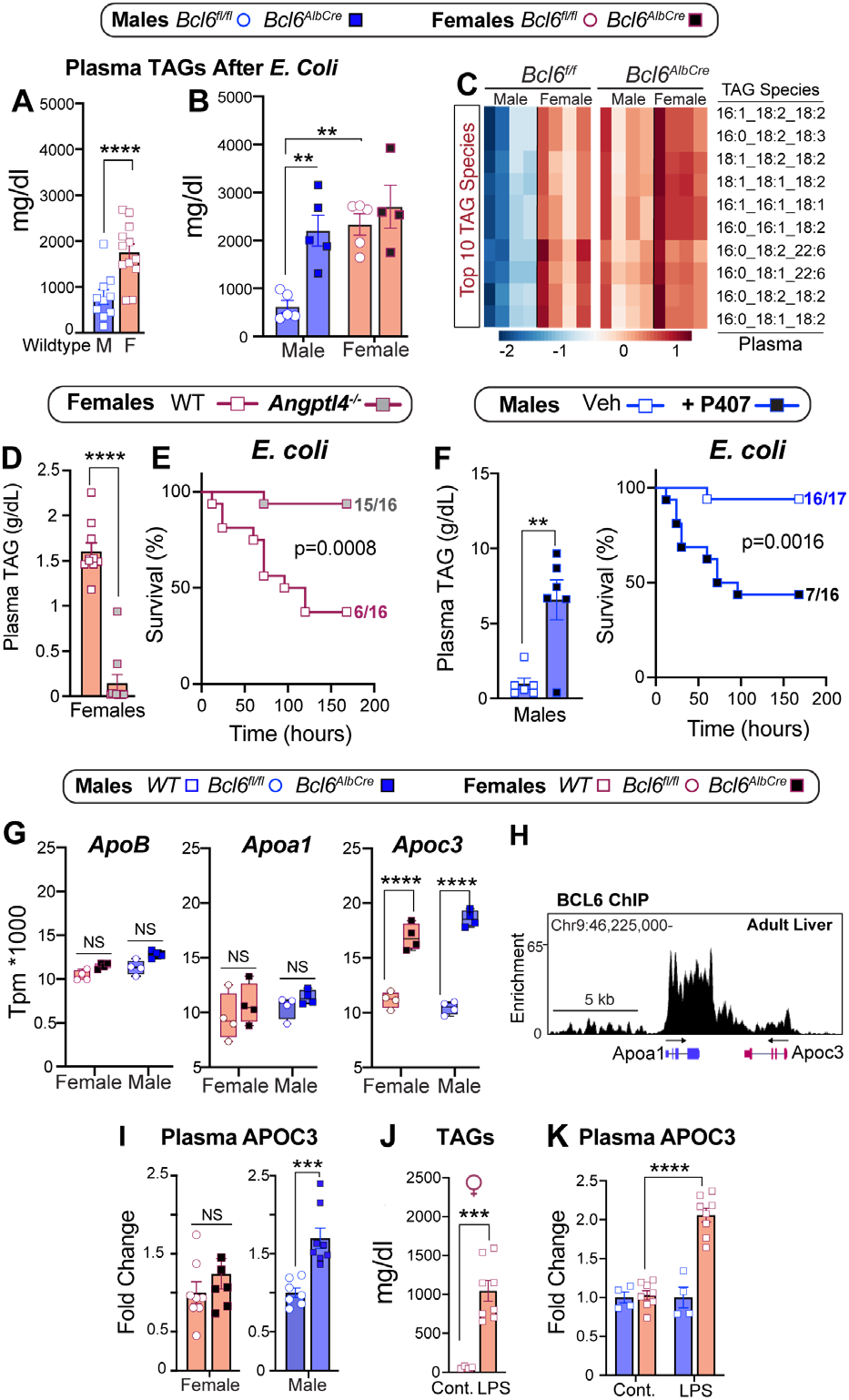
Sex-dependent Hepatic Hyperlipidemia and Lipids Profile Are Linked to Host Defense Responses. **(A)-(D)** Plasma triglycerides of mice infected with *E. coli.* **(E)** survival curves of mice infected with E. coli (1×10^8^ CFU) pooled from 2 experiments. **(F)** Plasma triglycerides and survival curves of mice infected with E. coli (1×10^8^ CFU). **(G)** Transcript abundance of hepatic *ApoB, Apoal,* and *Apoc3* in *Bcl6^f/f^* and *Bcl6^AlbCre^* mice. **(H)** c binding of BCL6 in *Apoc3/Apoal* locus in the male liver (ChIPseq). **(I)** Plasma APOC3 in *Bcl6^f/f^* and *Bcl6^AlbCre^* mice housed at 30°C. **(J-K)** Plasma triglycerides and APOC3 in control and LPS-treated (0.5 mg/kg) wildtype mice. Data are presented as mean ± SEM, NS: not significant. **p<0.01; ***p<0.001; ****p < 0.0001.

Increased plasma TAGs in infected *Bcl6^AlbCre^* mutant male mice prompted us to investigate whether crucial genes in packaging and clearance of very-low-density lipoprotein (VLDL) and their TAG cargo are regulated by BCL6. Of the three candidate genes examined, hepatic *Apoc3* expression increased dramatically in *Bcl6^AlbCre^* mice while *ApoB* and *ApoAl* were unchanged (Fig. 3G). Reanalysis of the hepatic BCL6 ChIP-Seq dataset by Waxman’s group revealed that BCL6 binds directly to the *Apoc3* locus to dampen its expression (Fig. 3H) (15). As predicted, eliminating high hepatic BCL6 in males increased circulating APOC3 (Fig. 3I). Given the essential role of hepatic APOC3 in maintaining circulating TAGs (18, 19), we suggest that high plasma TAGs in LPS-treated wildtype females (Fig. 3J) arise from high APOC3 and low BCL6 in females but not in males (Fig. 3K). How these changes in lipid handling alter host fitness during infection remains to be established.

We next asked when and what factors enable BCL6 to control the hepatic gene programs in male mice. The appearance of male-biased genes coincides with puberty, becoming apparent at eight weeks of age (Fig. 4A and fig. 4SA). Surgical castration (GDX) of pre-pubescent males enhanced female-biased gene expression in the liver (fig. S4B), led to a steep drop in survival accompanied by elevated plasma TAGs (fig. S4C, D), and diminished hepatic BCL6 levels, which were partially restored by Testosterone (T) treatment (fig. S4E). Pulsatile secretion of growth hormone (GH) from the anterior pituitary is distinct in males and consists of peaks with prolonged extended dips – this unique pattern is required for male-biased hepatic gene expression in mice (15, 20). As predicted from (21), continuous infusion of GH feminizes male livers (Fig. 4B and fig. S4F) and results in dramatic repression of hepatic BCL6 protein and transcripts (Fig. 4C). While saturating levels of GH in primary hepatocytes also suppressed *Bcl6*, T and estradiol (E) had no effect in this setting, implying that the T-induced rescue of BCL6 expression in vivo must be indirect (Fig. 4D and fig. S4E). Expectedly, disrupting normal GH pulsatility by continuous GH infusion reduced hepatic BCL6 and diminished host fitness in males (Fig. 4E). The major effector of hepatic GH signaling, STAT5, binds to the *Bcl6* locus (15), providing a direct molecular link between GH and BCL6 levels (Fig. 4F).

**Figure 4.**
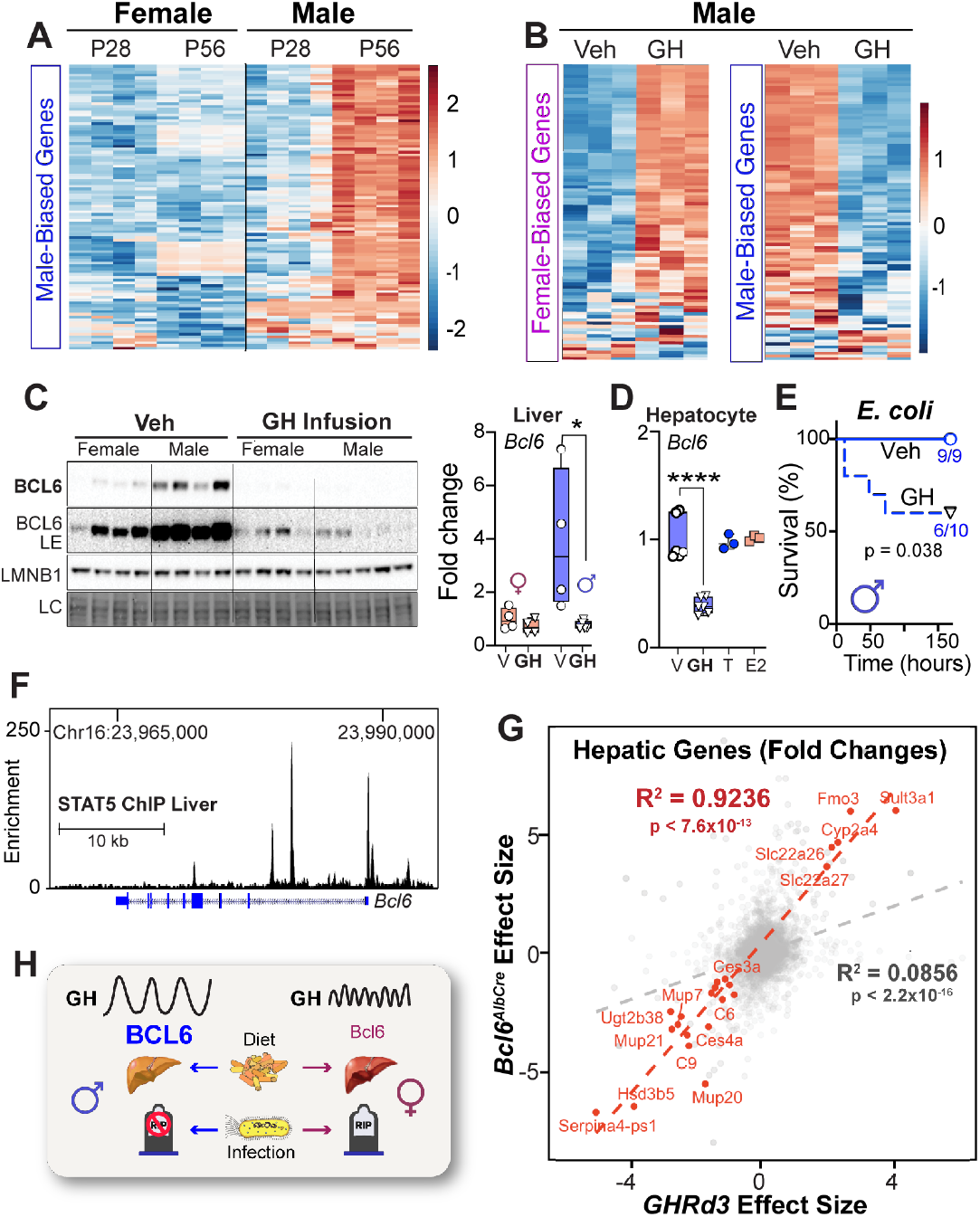
Sex-dependent GH Signaling Controls BCL6 Levels and Survival to Infection. Heatmaps of top 100 **(A)** male-biased genes at P28/P56 and **(B)** female/male-biased genes after GH treatment at 22°C, filtered by fold-change. **(C)** Immunoblotting and RT-QPCR for hepatic BCL6 protein and transcript in mice infused with vehicle (Veh) or recombinant mouse growth hormone (GH) for 15 days at 22°C. **(D)** *Bcl6* mRNA levels in primary mouse hepatocytes treated with vehicle (V), testosterone propionate (T), estradiol benzoate (E2) (n=3) or GH (n=6). **(E)** Survival curves of male mice infused with Veh or GH for 13 days and then infection with *E. coli* (1×10^8^ CFU) at 30°C (n=10 per group). **(F)** STAT5 binding to *Bcl6* locus in the male liver (ChIPseq). **(G)** Effect size correlation of all (gray) or differentially expressed (red) transcripts in livers of *Bcl6^AlbCre^* and *Ghrd3* male mice. **(H)** Schematic of GH-BCL6 axis in regulating sex-dependent endpoints when challenged by diet or infection. Data are presented as mean ± SEM, *p<0.05; ****p < 0.0001.

To extend these findings, we leveraged a mouse model carrying the common human variant of Growth Hormone Receptor *(GHRd3,* deletion of exon 3) that mimics increased GH signaling. This variant feminizes livers in male mice and is thought to confer an evolutionary advantage during periods of food scarcity in humans (22). The significant overlap in hepatic gene changes detected in *Ghrd3* and *Bcl6A^lbCre^* mutant males and the trend towards lowered *Bcl6* in *Ghrd3* male livers (Fig. 4G and fig. S4F) suggest that this common *GHR* variant attenuates hepatic BCL6 function. Further, the ability of the *GHRd3* variant to stave off nutritional stress (22), coupled with our study, implies that the GH-BCL6 signaling axis creates a trade-off for females that diminishes fitness during infection while enhancing survival in the fasted state. Indeed, in most recorded famines, survival rates for women outpace men (23).

Although the male-biased hepatic GHR-BCL6 axis is essential for mounting protective defenses against pathogens, this pathway leads to substantial hepatic fat accumulation and glucose intolerance during caloric excess (Fig. 4H). Thus, the current prevalence of fatty liver in men might stem from older host defense mechanisms postulated to have coevolved with sexually dimorphic aggressive behaviors required for mating and social status (24).

In summary, we propose that the strikingly different responses in male and female mice to an infectious or dietary challenge offer new evidence that modern sex-biased diseases are rooted in ancient evolutionary trade-offs between immunity and metabolism.

## METHODS

### Animals

Animal studies were conducted under an approved Institutional Animal Care and Use Committee (IACUC) protocol at the University of California, San Francisco (UCSF). Male and female mice were housed under a 12-hr light:dark cycle with free access to food and water. Unless otherwise indicated all studies were performed with C57BL/6J (Stock No: 000664) 8 to 10-week-old male and female mice housed at 30°C in a laboratory incubator (Darwin Chambers) for 2 weeks prior to initiation of studies. *Bcl6^f/f^* (Stock No. 023727) and Alb-Cre (Stock No. 003574) were purchased from Jackson Laboratories and bred to generate appropriate alleles; all alleles were backcrossed onto the C57BL6/J background for >10 generations. *Angptl4^-/-^* (Angptl4tm1Jig) mice were kindly provided by Dr. Koliwad (UCSF). For in vivo studies, cohorts of greater than or equal to 3 mice per genotype, sex, and treatment were assembled, and experiments were repeated 2-3 times.

### E. coli infection and LPS studies

*Escherichia coli* (strain O111:B4) was obtained from the laboratory of Dr. Judith Hellman (UCSF). *E. coli* was grown to log phase at an OD660 value of 0.8 in LB Broth (Lennox formulation, Sigma), collected by centrifugation at 4000g, washed once, and resuspended in fresh LB broth. *E. coli* was grown in large batches and stored in small aliquots at −80°C. An aliquot of *E. coli* was thawed once, diluted in phosphate-buffered saline (PBS), and plated on LB agar to determine the colony-forming units (CFU) before use in infection studies. The LD50 of each bacterial batch was tested by infecting C57BL/6J mice at thermoneutrality. *E. coli* batches with similar LD50 were used.

For *E. coli* survival studies, mice were infected at indicated CFU of *E. coli*, and body mass was recorded at the indicated times. Mice were monitored for absolute survival and were euthanized if they lost >25% of initial body mass or were found unresponsive in the cage. To quantify bacterial burden in tissues, spleen was collected from euthanized mice 24 hours after infection with *E. coli.* Tissues were homogenized in ice-cold PBS (200-400 mg/ ml) and lysates were plated on LB agar at 1:1-1:1,000 dilution for quantification of tissue bacterial burden. Lipopolysaccharides purified from *E. coli* O111:B4 (Sigma-Aldrich, #L3024) were diluted in sterile saline and injected at 1.75-2 mg/kg for survival experiments, as indicated in the figure legends. Mice were monitored undisturbed for absolute survival. For inhibition of lipoprotein lipase (LPL), poloxamer-407 (P-407; Sigma, P2443) was dissolved in ice-cold saline (250 mg/kg) and incubated in cold-room overnight on a rotating shaker. P-407 solution or saline (200 μl) was injected intraperitoneally into mice 1-2 hours before infection with *E. coli* and then daily until day seven or completion of the experiment.

### High-Fat Diet and GTT

High-fat diet was purchased from Research Diets (D12492), while standard diet was obtained from Pico Lab (5053). Glucose tolerance tests were conducted after a 6 hour fast with glucose administered intraperitoneally (1.5 g/(kg of lean mass)). Body composition was analyzed by dual-energy x-ray (DEXA, GE Lunar PIXImus).

### ISH and Liver Histology

Fluorescent in situ hybridization was performed using RNAScope(ACD; Multiplex Fluorescent v.2) according to the manufacturer’s instructions *(Bcl6* probe: 455311-C2). Lipid droplets were visualized in frozen liver sections with Oil Red O reagent (Millipore-Sigma; O1391) according to the manufacturer’s instructions. Slides were imaged with Keyence BZ-X800 fluorescence microscope. Three representative views of each liver sample were selected, and median droplet size was assessed with BZ-X800 Analyzer. H&E staining was done as service in Histology and Light Microscopy core at Gladstone Institutes.

### Gonadectomy and GH Infusion Studies

C57BL/5J male mice (4-5 weeks old) were gonadectomized (GDX) according to standard operating procedures under isoflurane anesthesia. Control mice underwent sham surgeries during which the gonads were exteriorized and then returned to the body cavity. Peri- and postoperative pain was managed using a combination of buprenorphine, meloxicam, and bupivacaine. GDX and sham mice were allowed to recover for at least two weeks before the initiation of experiments. Testosterone propionate USP (Spectrum Chemicals, T1315) was continuously infused via an osmotic pump (Alzet, pump model 1004) at a rate of 1.875 μg/hour, which yields physiological levels of circulating testosterone (25). Control mice received pumps filled with vehicle (ethanol in PEG-400). Pumps were implanted subcutaneously above and caudal to the scapulae at five weeks of age. Mice were adapted to thermoneutrality (30°C) at six weeks of age and infected with *E. coli.* at eight weeks of age. Survival and tissue collection were performed as described above.

Osmotic pumps (Alzet, pump model 1004) were implanted subcutaneously into mice to continuously deliver recombinant mouse growth hormone (obtained from A.F Parlow, National Hormone and Peptide Program, UCLA) at a rate of 15 ng/g/h in buffer containing 30 mM NaHCO3, 150 mM NaCl, 0.1 mg/ml rat albumin (Sigma Aldrich #A6414), pH 10.5. Growth hormone was infused into mice for 13-15 days before the initiation of experiments. Control animals received infusion at the same rate as the vehicle.

### Plasma and liver TAG and APOC3 Assays

Triglycerides (Cayman Chemicals) in plasma and tissues were quantified using commercially available kits, as per manufacturer’s protocols. All plasma and liver samples were stored at −80°C prior to analysis. APOC3 plasma levels were measured using ELISA assay (Abcam #AB217777).

### Immunoblotting

Whole-cell extracts were prepared in buffer containing 50 mM Tris-HCl pH 7.5, 150 mM NaCl, 0.1% w/v SDS, 0.5% w/v Na-Deoxycholate, 1% v/v Nonidet P-40 substitute (Fluka, #74385), 1mM EDTA, 1mM EGTA and protease inhibitor cocktail (Sigma-Aldrich, #P8340). To detect BCL6 expression in liver samples, nuclear extracts were prepared using a nuclear extraction kit (Thermo Fisher, #78833) according to the manufacturer’s instructions. The protein samples were run into Criterion or mini-PROTEAN TGX Stain-Free precast gels (Biorad) and blotted onto a nitrocellulose membrane. The membranes were blocked (5% non-fat milk in 0.1% TBST) for one h at room temperature and probed with anti-BCL6 antibody (Cell Signaling, #5650S) and anti-Lamin B (Cell Signaling, #12586. dilution 1:1000) antibody in 0.1% TBST containing 3% BSA overnight at +4°C followed by incubation with secondary HRP-linked anti-rabbit antibody (Cell Signaling, #7074). Immunoblotted proteins were detected with ProSignal Pico ECL Reagent (Genesee Scientific, #20-300) and imaged using ChemiDoc MP Imaging System (Bio-Rad). *Primary Hepatocyte Cultures*

Primary hepatocytes were isolated from C57BL/6J male mice by perfusing liver through superior vena cava with liver perfusion medium (Thermo Fisher Scientific, #17701038) and liver digest medium (Thermo Fisher Scientific, # 17703034). Live hepatocytes were purified with Percoll reagent (Sigma Aldrich, #GE17-0891-01) and plated on Collagen I coated cell culture plates (Thermo Scientific, #12-565-907) in hepatocyte plating medium (DME H21 high glucose medium without phenol-red (UCSF), 1 x penicillinstreptomycin (UCSF), 1 x insulin-transferrin-selenium solution (Gibco #41400045) and 5 % (v/v) charcoal-stripped fetal bovine serum (Thermo Fisher Scientific, #A3382101)). Hepatocytes were cultured 24 h before treatment with 300 ng/ml recombinant mouse growth hormone (A.F Parlow, National Hormone and Peptide Program, UCLA), 10 nM testosterone propionate (Spectrum Chemicals, #T1315), or 10 nM estradiol benzoate (Cayman Chemicals, #10006487) for five hours before analysis.

### Quantitative RT-PCR

Total RNA was used to synthesize cDNA (QuantaBio, #95048), and quantitative PCR was performed using SensiFAST SYBR reagent (Bioline, #BIO-98020). Transcript abundance was normalized using Ywhae as a reference gene. Primer sequences used are as follows:

*Bcl6:* Fwd 5’-GACGCACAGTGACAAACCAT-3’ Rev 5’-GATTGAACTGCGCTCCACAA-3’
*Ywhae:* Fwd 5’-ATGCAGGGTGATGGTGAAGAG-3’ Rev 5’-TGTTGGCTTTTATTTCGTCTCAC-3’

### Chromatin Immunoprecipitation

Snap-frozen liver samples were minced and fixed with 1% formalin in PBS for 15 minutes at room temperature. The samples were treated with glycine (final concentration 444 mM), washed with PBS, and collected in cell lysis buffer (50 mM Tris-Cl, pH 8.0, 140 mM NaCl, 1mM EDTA, 10% glycerol, 0.5% Nonidet P-40 substitute (Fluka, #74385), 0.25% Triton X-100 and protease inhibitor cocktail (Sigma-Aldrich, #P8340)). Nuclei were collected and lysed in nuclear lysis buffer (10 mM Tris-Cl, pH 8.0, 1 mM EDTA, 0.5 mM EGTA, 0.2% SDS and protease inhibitor cocktail). Extracted chromatin was sheared by sonicating the samples with an XL-2000 sonicator (QSonica) for 12 rounds on setting 2. The samples were incubated with anti-H3K27ac antibody (Abcam, #ab4729) overnight at +4°C and captured with Protein G Dynabeads for 4 hours at 4°C.

### Next-Generation Sequencing Library Preparation

RNA sequencing libraries were prepared using the TruSeq Stranded mRNA library prep kit (Illumina) according to instructions. ChIP DNA was first treated with end repair enzymes (Thermo Scientific, #K0771) before continuing to library preparation according to Illumina’s instructions. The pooled libraries were sequenced on NextSeq 500 Illumina sequencer using NextSeq 500/550 High Output Kit v2.5 (75 cycles) (Illumina #20024906) in a single read mode.

### RNA-seq and ChIP-seq Analysis

Sequences were aligned to mouse transcriptome (mm10) using Kallisto in gene mode (26). Differential gene expression was evaluated using the likelihood-ratio test by Sleuth (qval <0.05) (27). All heatmaps were generated with top 100 female/male-biased genes obtained from 8-week-old mice housed at 22°C (Fig. 2C, D; listed in Data S1). Principle Components Analysis (PCA) plots and heat maps were generated in R (28). Sequences from ChIP experiments were aligned to the mouse genome (mm10) using Bowtie2 (29). Subsequent analysis to identify and annotate peaks was done with Homer (30). DESeq2 was used to evaluate differential enrichment of (31) and DeepTools was used to create heatmaps of differentially enriched peaks (32). Sex-biased H3K27ac peaks were identified comparing female (n=3) and male (n=3) samples using Homer command getDifferentialPeaksReplicates in histone mode (qval <0.05).

### Lipidomics

Lipidomics was performed at the Metabolomics Core Facility at the University of Utah. Plasma samples were processed for untargeted lipidomics as described. Extraction protocols were based on (33). All solutions are pre-chilled on ice. Tissues are transferred to labeled bead-mill tubes (1.4 mm, MoBio Cat# 13113-50) where lipids are extracted in a solution of 250 μL PBS, 225 μL MeOH containing internal standards (Avanti SPLASH LipidoMix (Lot#12) at 10 μL per sample; Cambridge Isotope laboratories NSK-B and NSK-B-G1 (deuterated carnitines) at 10 μL per sample) and 750 μL MTBE (methyl tert-butyl ether). The sample is homogenized in one 30 s cycle using the Omni Bead Ruptor followed by a rest on ice for 1 hour. Plasma samples are transferred (25 uL) to glass vials containing the same extraction solution described above but without PBS. Samples are sonicated for 1minute, then rest on ice for 1 hour, after which an addition of 188 μL PBS is made to induce phase separation. After centrifugation at 16,000 g for 5 minutes at 4 °C, the upper phases are collected and evaporated to dryness under a gentle nitrogen stream at room temperature. Lipid samples are reconstituted in 500 μL IPA and transferred to an LC-MS vial with insert (Agilent 5182-0554 and 5183-2086) for analysis. Concurrently, a process blank sample and pooled quality control (QC) sample is prepared by taking equal volumes (~25 μL) from each sample after final resuspension.

### Mass Spectrometry Analysis of Samples

Lipid extracts are separated on a Waters Acquity UPLC CSH C18 1.7 μm 2.1 x 100 mm column maintained at 65 °C connected to an Agilent HiP 1290 Sampler, Agilent 1290 Infinity pump, equipped with an Agilent 1290 Flex Cube and Agilent 6530 Accurate Mass Q-TOF dual AJS-ESI mass spectrometer. For positive mode, the source gas temperature is set to 225 °C, with a drying gas flow of 11 L/min, nebulizer pressure of 40 psig, sheath gas temp of 350 °C and sheath gas flow of 11 L/min. VCap voltage is set at 3500 V, nozzle voltage 1000V, fragmentor at 110 V, skimmer at 85 V and octopole RF peak at 750 V. For negative mode, the source gas temperature is set to 300 °C, with a drying gas flow of 11 L/min, a nebulizer pressure of 30 psig, sheath gas temp of 350 °C and sheath gas flow 11 L/min. VCap voltage is set at 3500 V, nozzle voltage 2000V, fragmentor at 100 V, skimmer at 65 V and octopole RF peak at 750 V. Samples are analyzed in a randomized order in both positive and negative ionization modes in separate experiments and acquired with the scan range m/z 100 – 1700. Mobile phase A consists of ACN:H2O (60:40 v/v) in 10 mM ammonium formate and 0.1% formic acid, and mobile phase B consists of IPA:ACN:H2O (90:9:1 v/v) in 10 mM ammonium formate and 0.1% formic acid. The chromatography gradient for both positive and negative modes starts at 15% mobile phase B then increases to 30% B over 2.4 min, it then increases to 48% B from 2.4 – 3.0 min, then increases to 82% B from 3 – 13.2 min, then increases to 99% B from 13.2 – 13.8 min where it’s help until 16.7 min and then returned to the initial conditioned and equilibrated for 5 min. Flow is 0.4 mL/min throughout, injection volume is 2 μL for positive and 10 μL negative mode. Tandem mass spectrometry is conducted using the same LC gradient at collision energy of 25 V.

### Analysis of Mass Spectrometry Data

QC samples (n=8) and blanks (n=4) are injected throughout the sample queue and ensure the reliability of acquired lipidomics data. Results from LC-MS experiments are collected using Agilent Mass Hunter (MH) Workstation and analyzed using the software packages MH Qual, MH Quant, and Lipid Annotator (Agilent Technologies, Inc.). Results from the positive and negative ionization modes from Lipid Annotator are merged then split based on the class of lipid identified. For example, we detect PC in both POS and NEG modes, but only use the NEG mode data to tabulate data for those targets. Lipid targets are normalized to the spiked internal standards in MH Quant. The data table exported from MHQuant is evaluated using Excel where initial lipid targets are parsed based on the following criteria. Only lipids with relative standard deviations (RSD) less than 30% in QC samples and are used for data analysis. Additionally, only lipids with background AUC counts in process blanks that are less than 30% of QC are used for data analysis. The parsed excel data tables are normalized to tissue mass and positive and negative mode data are merged.

### Data availability

RNA-seq and ChIP-seq datasets have been deposited at GEO (https://www.ncbi.nlm.nih.gov/geo/) under the SuperSeries accession number GEO: GSE138396. RNAseq analysis from livers of male mice infused with growth hormone was performed using GEO samples SRX2786690-SRX2786694 and SRX883050-SRX883061. ChIP-seq dataset for STAT5 (GSE31578) was used to identify STAT5 binding sites across the Bcl6 gene in livers of male and female mice.

### Statistical Analysis

Statistical analysis was performed using Prism 8 (GraphPad Software, San Diego, CA, USA). Data are presented as mean ± s.e.m. Statistical significance was determined using the unpaired two-tailed Student’s t-test for single variables, one-way ANOVA for multiple cohorts, and two-way ANOVA followed by Bonferroni post-tests for multiple variables. For survival experiments, statistical significance was determined using the Mantel-Cox log-rank test with a p value of < 0.05. For all figures, *p < 0.05, **p < 0.01, ***p < 0.001 and ****p<0.0001. Biological replicates for each of the studies are indicated in scatter plots and also listed in each figure legend.

## ACKNOWLEDGMENTS

We thank members of the Chawla and Ingraham labs for comments on the manuscript, and X. Cui and J. Argiris for assistance with mouse husbandry. The authors’ work was supported by grants from NIH (DK094641, DK101064) and the Pathology & Imaging Core of the UCSF Liver Center (P30 DK026743). Y.A.L. was supported by NHMRC (GNT1142229), and J.N. by EMBO (ALTF 1185-2017), HFSPO (LT000446/2018-L), and PBBR (7000/7002124). Mass spectrometry equipment was obtained through NIH Shared Instrumentation Grant 1S10OD016232-01 (J.E.C), 1S10OD018210-01A1 (J.E.C), and 1S10OD021505-01 (J.E.C). The authors declare that they have no competing financial interests.

## AUTHOR CONTRIBUTIONS

J.N., Y.A.L., W.C.K., K.C.C, K.G., E.J.W., H.A.I, and A.C. conceived and designed the experiments, interpreted the results, and wrote the paper. J.N., Y.A.L., W.C.K., K.C.C, and J.L.T. performed the experiments. D.D. assisted with analysis of RNA-seq and ChIP-seq datasets, and J.A.M, T.V.R, J.E.C. performed lipidomics on plasma and liver samples.performed analyses, and wrote the manuscript. H.A.I designed experiments, analyzed data, and wrote the manuscript.

Supplementary materials will be provided upon request.

